# Examining the impact of the *Chlamydia muridarum*-induced synthesis of IFN-β during genital tract infection^1^

**DOI:** 10.64898/2026.03.23.713660

**Authors:** Ramesh Kumar, Iliana C. Cordova-Mendez, Fnu Litika, Evan D. Kara, Rayna Moiz, Derrick Burgess, Arkaprabha Banerjee, Wilbert A. Derbigny

**Affiliations:** Department of Microbiology, Marian University-Wood College of Osteopathic Medicine Indianapolis, Indiana 46222; Department of Pediatrics, Indiana University School of Medicine Indianapolis, Indiana 46202; Dept. of Microbiology; Indiana University School of Medicine Indianapolis, Indiana 46202; Department of Neuroscience, Butler University Indianapolis, Indiana 46268

**Keywords:** Oviduct epithelial cells, *Chlamydia*, IFN-β-deficiency, cytokines, epithelial, mucosa, inclusion bodies

## Abstract

*Chlamydia trachomatis* infection of the female genital tract can result in severe reproductive sequelae, including pelvic inflammatory disease, tubal scarring, and infertility. Type I interferons have been implicated in both host defense and immunopathogenesis during chlamydial infection, with conflicting conclusions across experimental systems. However, the specific contributions of individual interferon subtypes remain poorly defined. Here, we examined the role of interferon beta (IFN-β) in regulating epithelial immune responses and intracellular bacterial development during *Chlamydia muridarum* infection.

Using murine oviduct epithelial (OE) cell lines derived from wild-type, IFNβ–deficient, and Toll-like receptor 3 (TLR3)–deficient mice, we demonstrate that IFN-β is a critical epithelial-intrinsic mediator of host defense. Loss of IFN-β led to dysregulation of genes associated with inflammation, immune regulation, and fibrosis, altered chlamydial inclusion morphology, enhanced expression of bacterial genes throughout the chlamydial developmental cycle, and increased chlamydial replication. Importantly, exogenous IFN-β restored both immune mediator production and bacterial control during IFNβ-deficiency. Parallel analyses revealed that TLR3 deficiency phenocopied IFN-β loss, supporting a TLR3–IFN-β signaling axis that restricts chlamydial growth.

Consistent with these *in vitro* findings, IFNβ-deficient mice exhibited enhanced bacterial burden during genital tract infection. Together, these data establish IFN-β as a protective epithelial mediator during chlamydial infection and demonstrate that type I interferon responses are not functionally uniform. Our findings provide a mechanistic framework to reconcile the protective role of IFN-β with reports of reduced pathology in interferon-α/β receptor-deficient models and highlight the importance of dissecting individual interferon pathways in chlamydial immunopathogenesis.

**Importance:** Genital tract infection with *Chlamydia trachomatis* remains a leading cause of preventable infertility worldwide. Although type I interferons are widely viewed as contributors to chlamydial pathology, most studies have examined global interferon signaling rather than the roles of individual interferon subtypes. In this study, we demonstrate that interferon beta (IFN-β) plays a protective, epithelial-intrinsic role during chlamydial infection by restricting bacterial development and shaping local immune responses. These findings challenge the prevailing view that type I interferons are uniformly detrimental in this setting and reveal that distinct interferon subtypes can exert opposing effects on host defense and disease outcome. By defining a TLR3–IFN-β signaling axis that limits chlamydial replication, this work advances our understanding of epithelial immunity in the female genital tract and has important implications for the design of targeted immunomodulatory strategies to prevent chlamydia-induced reproductive pathology.

## Introduction

*Chlamydia trachomatis* urogenital serovars are the most common cause of bacterial sexually transmitted infections worldwide and represent a major public health burden among women of reproductive age (1–3). Although many genital tract infections remain asymptomatic, persistent or recurrent *Chlamydia* infections can result in ascending infection of the upper female genital tract (FGT), leading to pelvic inflammatory disease, tubal scarring, hydrosalpinx, ectopic pregnancy, and infertility. Importantly, the immune responses required for pathogen clearance are also central drivers of tissue damage, underscoring the need to define immune pathways that balance host defense with preservation of reproductive tract integrity (1, 2, 4–8).

The innate immune response to *Chlamydia* infection is initiated predominantly by epithelial cells lining the FGT, which serve as both the primary targets of infection and critical sentinels of host defense. These epithelial cells express a diverse repertoire of pattern recognition receptors (PRRs) that detect microbial products and trigger the synthesis of inflammatory mediators, chemokines, and interferons that shape downstream immune responses (9, 10). While activation of innate signaling pathways is essential for controlling bacterial replication and recruiting immune effector cells, dysregulated or excessive epithelial responses can promote immunopathology and long-term tissue remodeling.

Type I interferons (IFNs), classically associated with antiviral immunity, have emerged as important modulators of host responses to intracellular bacterial pathogens, including *Chlamydia*. However, the role of type I IFNs in genital tract *Chlamydia* infection remains controversial. Several studies using interferon-α/β receptor–deficient (IFNAR^-/-^) mice have concluded that global type I IFN signaling exacerbates disease, promoting bacterial persistence and pathology. These findings have led to the prevailing view that type I IFNs are broadly detrimental during genital tract *Chlamydia* infection. Notably, IFNAR deficiency ablates signaling by all type I IFNs, precluding resolution of the contributions of individual interferons and obscuring potential cell-type–specific effects (11–13).

Our laboratory has previously demonstrated that interferon beta (IFN-β) is induced during *Chlamydia muridarum* infection of murine oviduct epithelial (OE) cells in a largely Toll-like receptor 3 (TLR3)–dependent manner (3, 9, 14). In these studies, TLR3 signaling was shown to regulate not only IFN-β synthesis but also the expression of multiple inflammatory mediators, and loss of TLR3 resulted in enhanced *Chlamydia* replication both in vitro and in vivo. Importantly, we further showed that exogenous IFN-β could restore defective cytokine responses and limit bacterial replication in TLR3-deficient OE cells, suggesting that IFN-β itself plays a protective role in epithelial host defense (15–17). These observations contrasted with conclusions drawn from IFNAR-deficient models and raised the possibility that IFN-β may function distinctively from other type I interferons during genital tract infection.

We therefore hypothesized that IFN-β is a non-redundant, epithelial-intrinsic mediator of host defense during *Chlamydia* infection and that its role cannot be inferred from studies that employ global disruption of type I IFN signaling. To directly test this hypothesis, we generated and characterized OE cells derived from IFNβ–deficient mice and examined their responses to *C. muridarum* infection. By comparing wild-type, IFNβ–deficient, and TLR3-deficient OE cells, we sought to define the specific contribution of IFN-β to epithelial immune signaling, bacterial control, and cellular pathology.

In the present study, we demonstrate that IFN-β deficiency profoundly alters epithelial immune responses to *Chlamydia*, resulting in impaired induction of key inflammatory mediators, enhanced bacterial replication, and striking differences in chlamydial inclusion morphology. Moreover, we show that exogenous IFN-β restores Chlamydia-induced cytokine levels and attenuates bacterial burden in IFNβ–deficient OE cells, recapitulating key features of TLR3-dependent host defense. Together, these findings establish IFN-β as a critical epithelial-derived mediator that limits *Chlamydia* replication and shapes innate immune responses in the FGT, providing mechanistic insight into the divergent outcomes observed in models of global type I IFN signaling deficiency.

## Results

### IFN-β deficiency disrupts Chlamydia-induced expression of inflammatory immune mediators in oviduct epithelial cells

Because epithelial cells are the primary targets of *Chlamydia* infection in the female genital tract (9, 10), we first examined the impact of IFN-β deficiency on the epithelial synthesis of mediators associated with the inflammatory immune response during *Chlamydia muridarum* infection. Wild-type and IFNβ–deficient murine oviduct epithelial (OE) cells were either mock-treated or infected with *C. muridarum*, and supernatants were collected at defined time points post-infection for cytokine and chemokine analysis by ELISA.

In wild-type OE cells, *C. muridarum* infection induced robust secretion of the inflammatory chemokines CCL5 and CXCL10, as well as the cytokines IL-6 and CXCL16 (Fig. 1A–D). The induction of CCL5 and CXCL10 is consistent with well-established roles for IFN-β in regulating interferon-stimulated genes and chemokines downstream of type I interferon signaling during intracellular infection (18). In contrast, the role of IFN-β in regulating IL-6 and CXCL16 expression during *Chlamydia* infection has been far less well defined.

**Fig 1.**
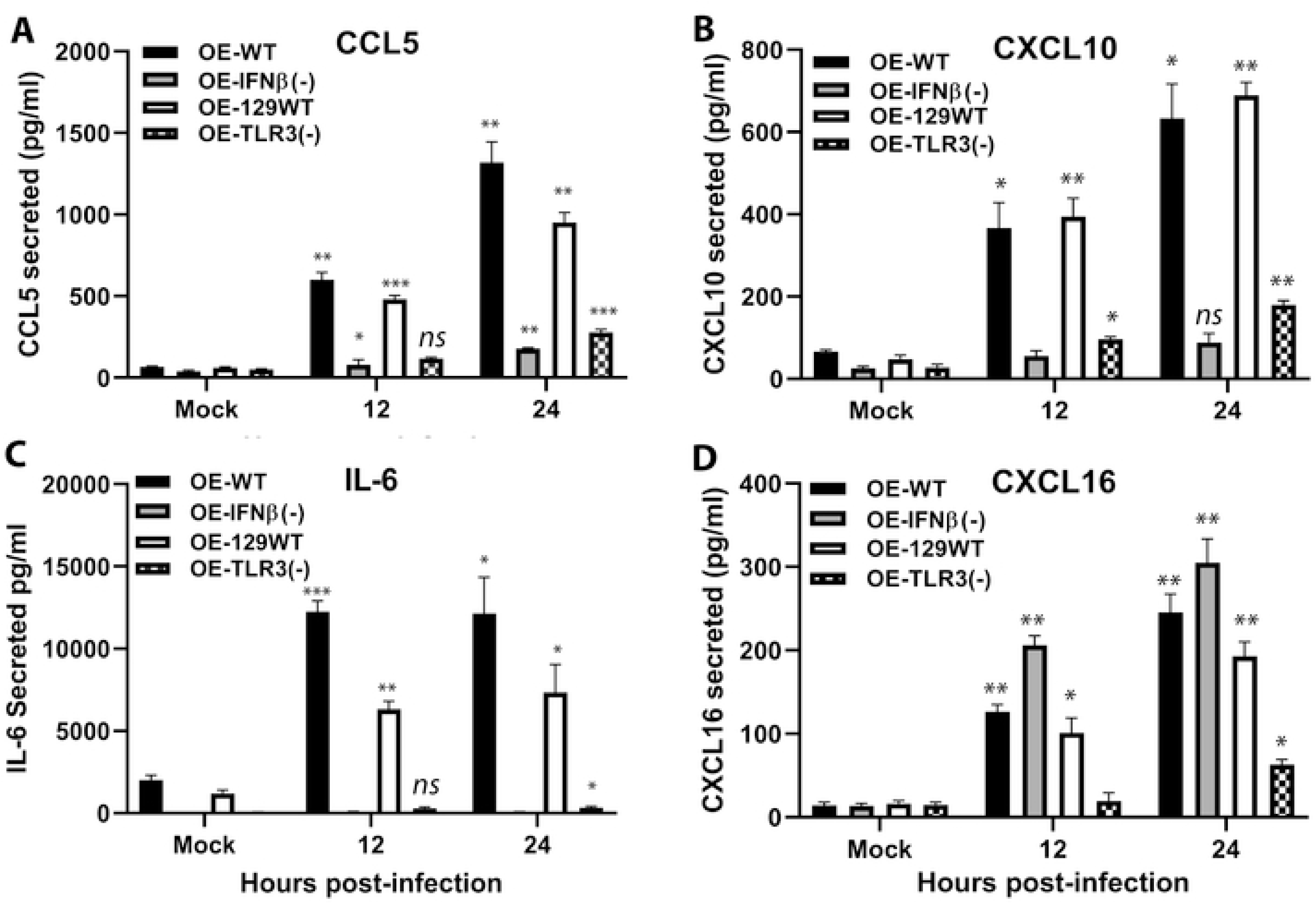
*Chlamydia*-induced expression of inflammatory cytokines in wild-type versus IFNβ-deficient OE cells. ELISA was used to measure Chlamydia-infection induced **(A)** CCL5, **(B)** CXCL10, **(C)** IL-6, and **(D)** CXCL16 secreted into the supernatants of wild-type and mutant OE cells that were either mock-treated or infected with the 10 IFU/cell *C. muridarum.* Supernatants were collected at the indicated time post-infection. *\*\*\*= p<0.005; **= p<0.01 *= p<0.05; ns= not statistically significant when compared to mock-infected control cells. Data presented are representative*.

Notably, IFNβ–deficient OE cells exhibited a marked reduction in Chlamydia-induced secretion of all four inflammatory mediators examined. While diminished CCL5 and CXCL10 production was anticipated based on their known association with IFN-β–dependent signaling pathways, the impaired inductions of IL-6 and CXCL16 in IFNβ–deficient OE cells were unexpected. These findings suggest that IFN-β contributes to epithelial inflammatory responses during *Chlamydia* infection through mechanisms that extend beyond classical interferon-stimulated gene regulation.

Together, these data demonstrate that IFN-β is required for optimal induction of a defined subset of inflammatory immune mediators in OE cells during *Chlamydia* infection and identify IL-6 and CXCL16 as previously underappreciated components of IFN-β–dependent epithelial immune signaling.

### Exogenous IFN-β restores inflammatory cytokine synthesis in IFNβ-deficient OE cells

To determine whether the impaired cytokine responses observed in IFNβ-deficient OE cells were directly attributable to the absence of IFN-β, we next assessed whether exogenous IFN-β could rescue *Chlamydia*-induced cytokine production. IFNβ-deficient OE cells were infected with *C. muridarum* in the presence or absence of recombinant IFN-β, and cytokine secretion was measured by ELISA.

Addition of exogenous IFN-β restored *Chlamydia*-induced IL-6 and CXCL16 secretion in IFNβ-deficient OE cells to levels comparable to those observed in infected wild-type cells (Fig. 2A–B). Treatment with recombinant IFN-β alone did not induce substantial cytokine production in the absence of infection, indicating that IFN-β acts synergistically with *Chlamydia*-induced signaling pathways rather than serving as a nonspecific inflammatory stimulus.

**Fig 2.**
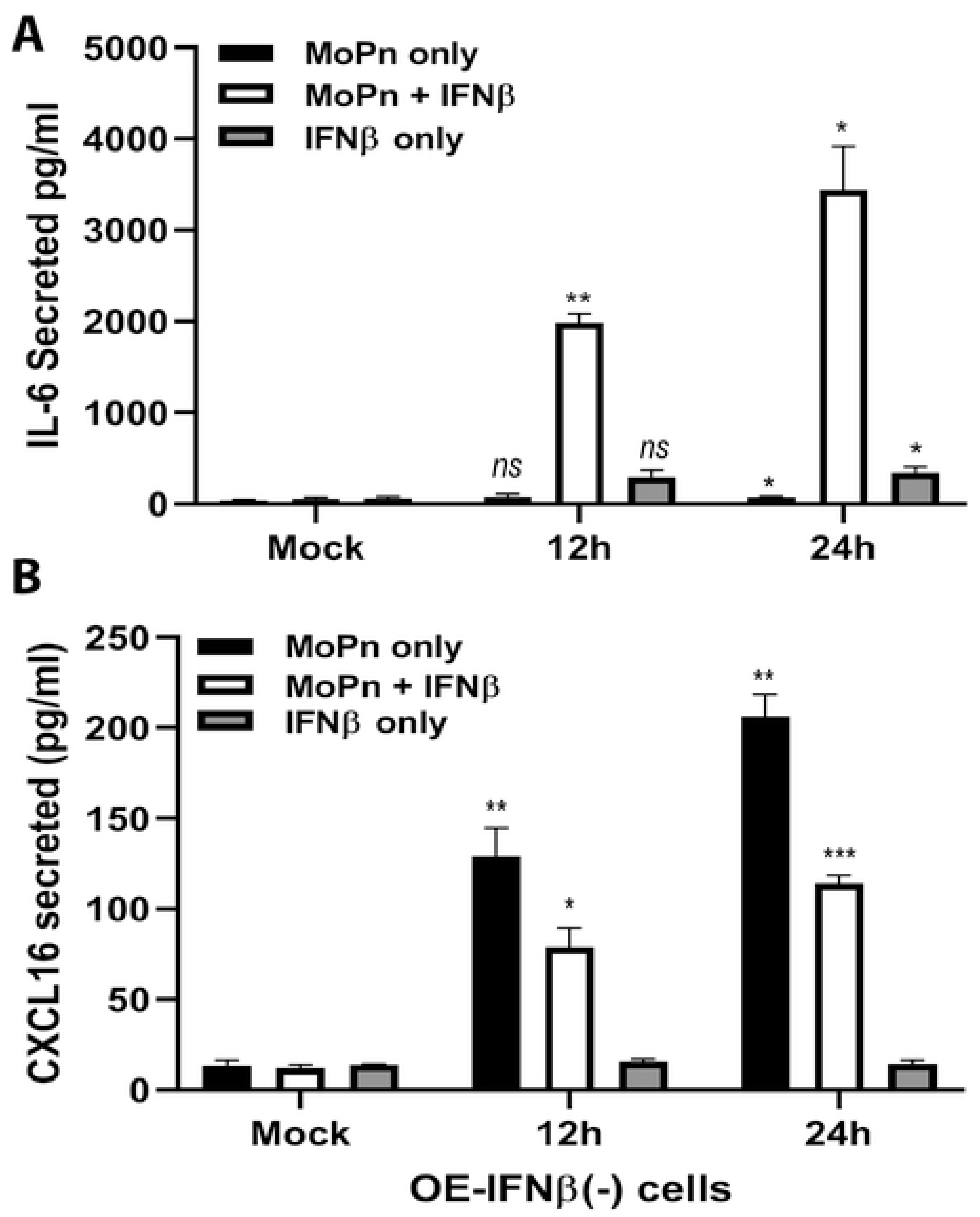
Exogenously added IFN-β restores the *Chlamydia*-induced expression of inflammatory cytokine synthesis in IFNβ-deficient OE cells. ELISA was used to measure *Chlamydia-*infection-induced **(A)** IL-6 and **(B)** CXCL16 secreted into the supernatants of OE-IFNβ(-) cells that were either mock-treated, infected with the 5 IFU/cell *C. muridarum* in the presence or absence of 50U of recombinant IFN-βml, or treated with only 50U of recombinant IFN-β/ml. Supernatants were collected at the indicated time post-treatment. *\*\*\*= p<0.005; **= p<0.01; *= p<0.05; ns= not statistically significant when compared to mock-infected control cells or where indicated. Data presented are representative*.

Together, these findings confirm that the defective cytokine responses observed in IFNβ-deficient OE cells are a direct consequence of IFN-β loss and demonstrate that IFN-β is both necessary and sufficient to restore some of these key components of the epithelial inflammatory response during *Chlamydia* infection.

### IFN-β deficiency alters epithelial gene expression profiles associated with immune regulation and fibrosis

Because dysregulated epithelial inflammatory responses are closely linked to tissue remodeling and pathology in the upper female genital tract (29, 30), we next examined whether IFN-β deficiency affects the transcriptional regulation of mediators involved in immune modulation and fibrosis during *Chlamydia muridarum* infection. Wild-type and IFNβ–deficient OE cells were either mock-treated or infected with *C. muridarum*, and gene expression was assessed by quantitative real-time PCR.

In wild-type OE cells, *Chlamydia* infection induced transcription of several cytokines and chemokines implicated in immune regulation and tissue remodeling (19), including IL-10, MMP9, IL-4, TNF-α, CCL5, and IFN-γ (Fig. 3A–F). In contrast, IFNβ–deficient OE cells displayed a markedly altered transcriptional profile following infection. Notably, the induction of MMP9 and IL-10, molecules associated with extracellular matrix remodeling and immunoregulatory balance (19), was significantly dysregulated in the absence of IFN-β. Similarly, transcription of TNF-α and IFN-γ, key mediators of inflammatory amplification and immune polarization, was altered in IFNβ–deficient cells compared with wild-type controls.

**Fig 3.**
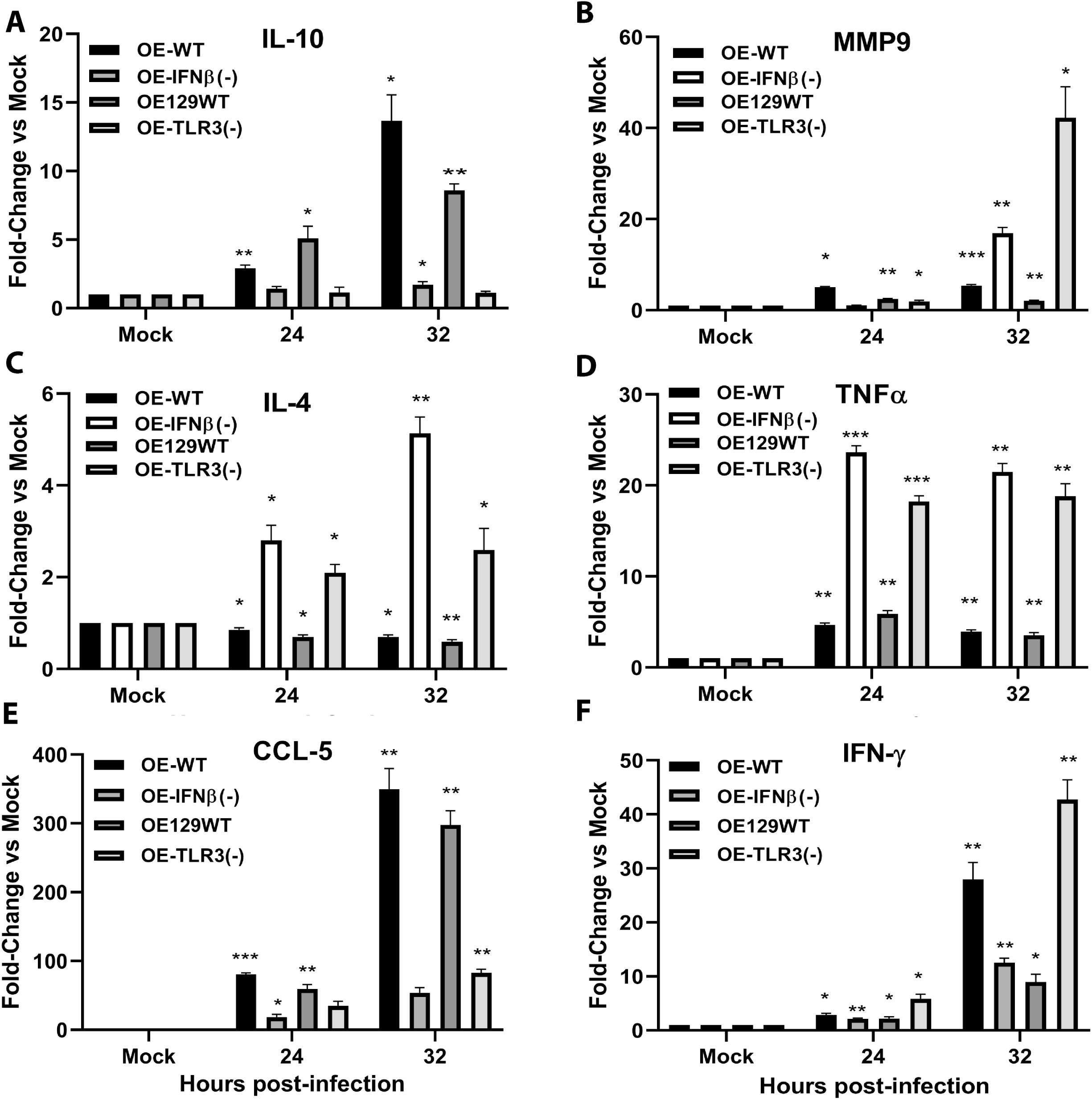
Impact of IFNβ-deficiency on the gene expression of cell mediators involved in fibrosis and scarring. Gene transcription levels of **(A)** IL-10, **(B)** MMP9, **(C)** IL-4, **(D)** TNFα, **(E)** CCL-5, and **(F)** IFN-γ were measured by real-time quantitative PCR (qPCR) in the various wild-type and gene-deficient OE cells that were either mock-treated or infected with 10 IFU/cell *Chlamydia muridarum* for the time indicated. Data are representative of 3 independent experiments. *\*= p <0.05; **= p <0.005; ***= p <0.001 compared to mock control*.

While several of these mediators have previously been linked to *Chlamydia*-induced inflammation and pathology, their dependence on epithelial-derived IFN-β has not been well defined. These findings suggest that IFN-β contributes to the coordinated regulation of immune and fibrosis-associated gene expression in OE cells during *Chlamydia* infection, providing a potential mechanistic link between epithelial innate signaling and downstream tissue remodeling processes.

### IFN-β deficiency results in altered chlamydial inclusion morphology in infected oviduct epithelial cells

Given the pronounced effects of IFN-β deficiency on epithelial immune signaling and gene expression, we next assessed whether loss of IFN-β impacted the intracellular developmental cycle of *Chlamydia* within OE cells. Wild-type and IFNβ–deficient OE cells were infected with *C. muridarum*, and chlamydial inclusions were visualized by immunofluorescence microscopy at 24 hours post-infection.

In wild-type OE cells, Chlamydia infection resulted in the formation of discrete, well-defined inclusions characteristic of normal chlamydial development (Fig. 4). In contrast, IFNβ–deficient OE cells exhibited striking differences in inclusion morphology, with inclusions appearing markedly larger and more prominent compared to those observed in wild-type cells. These morphological differences were consistently observed across independent experiments and were not present in mock-infected controls.

**Fig 4.**
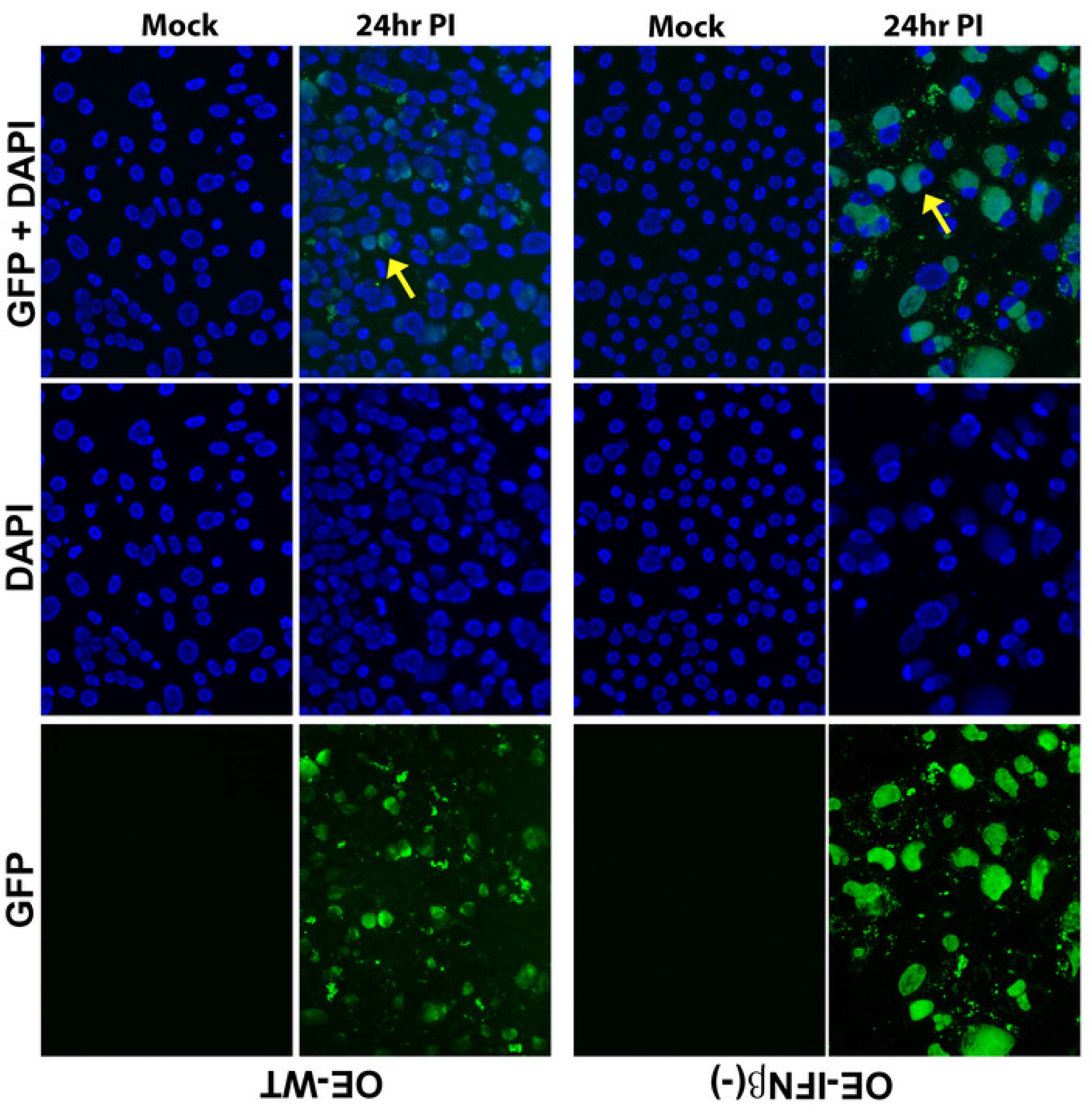
Microscopic analyses of chlamydial inclusions by immunofluorescence. Wild-type (OE-WT) and IFNβ-deficient (OE-IFNβ(-)) OE cells were either mock-infected or infected with *C. muridarum* at an MOI of 10 for 24 hours. The cells were stained with GFP-conjugated Chlamydia-specific monoclonal antibody (GFP), and nuclei were counter-stained with DAPI. The arrow indicates a typical chlamydial inclusion in each cell type. *Magnification=40X*

The altered inclusion morphology observed in IFNβ–deficient OE cells suggests that epithelial-derived IFN-β restricts *Chlamydia* growth or development at the cellular level. Together with the dysregulated immune gene expression profiles observed in IFNβ–deficient cells, these findings indicate that IFN-β plays a critical role in shaping both host epithelial responses and intracellular bacterial biology during *Chlamydia* infection.

### IFN-β deficiency enhances chlamydial gene expression during epithelial infection

Because IFN-β deficiency altered inclusion morphology and increased bacterial burden at the cellular level, we next examined whether loss of IFN-β directly affected chlamydial gene expression during infection of oviduct epithelial (OE) cells. Wild-type and IFNβ– deficient OE cells were either mock-treated or infected with *C. muridarum*, and transcription of representative chlamydial genes was quantified by real-time PCR (20, 21).

The chlamydial genes analyzed represent distinct stages of the chlamydial developmental cycle, including bacterial viability and transcriptional activity (16S rRNA), metabolic function (pyruvate kinase; *PyK*), structural components (outer membrane protein A; *OmpA*), and cell division (*FtsK*). In wild-type OE cells, *Chlamydia* infection resulted in time-dependent expression of each of these genes (Fig. 5A–D). In contrast, IFNβ–deficient OE cells exhibited significantly enhanced expression of PyK, 16S rRNA, OmpA, and FtsK at multiple time points post-infection. The increased transcription of genes associated with chlamydial metabolism, replication, and division is consistent with enhanced bacterial growth in the absence of IFN-β.

**Fig 5.**
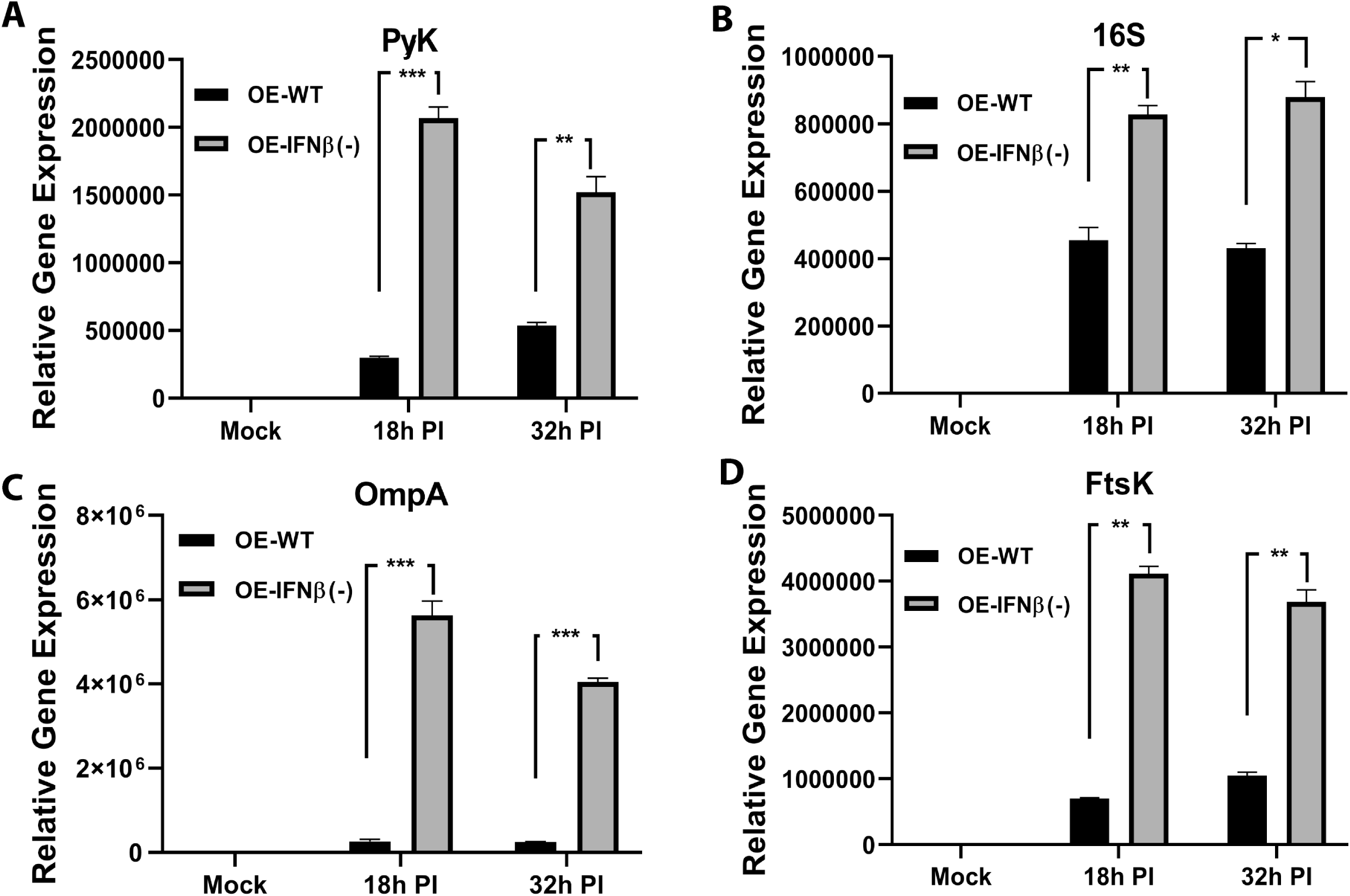
Impact of IFN-β deficiency on chlamydial gene expression. The chlamydial genes **(A)** PyK, **(B)** 16S, **(C)** OmpA, and **(D)** FtsK were measured by qPCR in wild-type and IFNβ-deficient OE cells that were either mock-treated or infected with 10 IFU/cell *Chlamydia muridarum* for the time indicated. Data are representative of 3 independent experiments*. *= p <0.05; **= p <0.005; ***= p <0.001 when comparing the cell lines at the respective time point*.

These findings demonstrate that epithelial-derived IFN-β restricts Chlamydia at the level of bacterial gene expression, supporting a role for IFN-β in limiting intracellular bacterial development.

### TLR3 deficiency phenocopies IFN-β deficiency with respect to chlamydial gene expression

Our previous studies established TLR3 as a critical upstream regulator of *Chlamydia*-induced IFN-β synthesis in OE cells. To determine whether the enhanced chlamydial gene expression observed in IFNβ–deficient OE cells was consistent with loss of TLR3 signaling, we performed parallel analyses using wild-type and TLR3-deficient OE cells infected with C*. muridarum*.

Similar to IFNβ–deficient OE cells, TLR3-deficient OE cells displayed significantly elevated expression of *PyK*, 16S rRNA, *OmpA*, and *FtsK* following infection compared to wild-type OE cells (Fig. 6A–D). The magnitude and kinetics of chlamydial gene expression in TLR3-deficient OE cells closely mirrored those observed in IFNβ–deficient OE cells, indicating that disruption of TLR3 signaling recapitulates the bacterial transcriptional phenotype associated with IFN-β loss.

**Fig 6.**
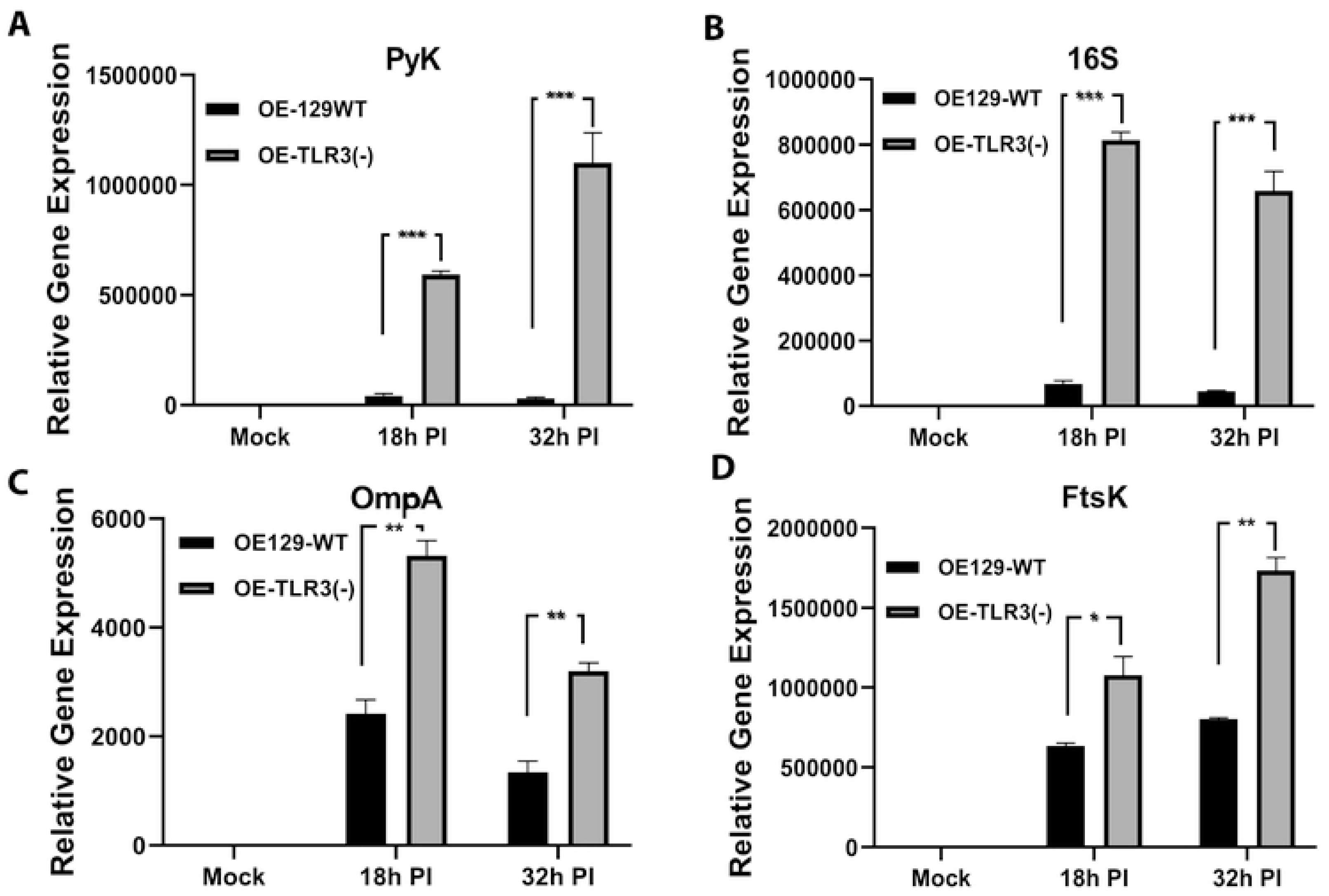
Impact of TLR3 deficiency on chlamydial gene expression. The chlamydial genes **(A)** PyK, **(B)** 16S, **(C)** OmpA, and **(D)** FtsK were measured by qPCR in wild-type and TLR3-deficient OE cells that were either mock-treated or infected with 10 IFU/cell *Chlamydia muridarum* for the time indicated. Data are representative of 3 independent experiments. *\*= p <0.05; **= p <0.005; ***= p <0.001 when comparing the cell lines at the respective time point*.

Together, these data support a model in which TLR3-mediated induction of IFN-β is a key epithelial signaling axis that limits chlamydial gene expression and intracellular replication during *Chlamydia* infection.

### IFN-β limits *Chlamydia muridarum* replication in oviduct epithelial cells

Because IFN-β deficiency enhanced chlamydial gene expression and altered inclusion morphology, we next determined whether IFN-β directly affected *Chlamydia* replication in OE cells. Wild-type and IFNβ–deficient OE cells were infected with *C. muridarum*, and bacterial replication was quantified by measuring recoverable inclusion-forming units (IFU) from infected cell lysates.

Consistent with the enhanced bacterial gene expression observed in IFNβ–deficient cells, loss of IFN-β resulted in significantly increased *C. muridarum* replication in OE cells compared to wild-type controls (Fig. 7). Importantly, the addition of exogenous recombinant IFN-β significantly attenuated *Chlamydia* replication in IFNβ– deficient OE cells, reducing bacterial titers to levels comparable to those observed in wild-type cells. In contrast, treatment with IFN-β had no significant effect on bacterial replication in wild-type OE cells, indicating that the observed restriction was specific to restoration of IFN-β–dependent signaling rather than a nonspecific antimicrobial effect.

**Fig 7.**
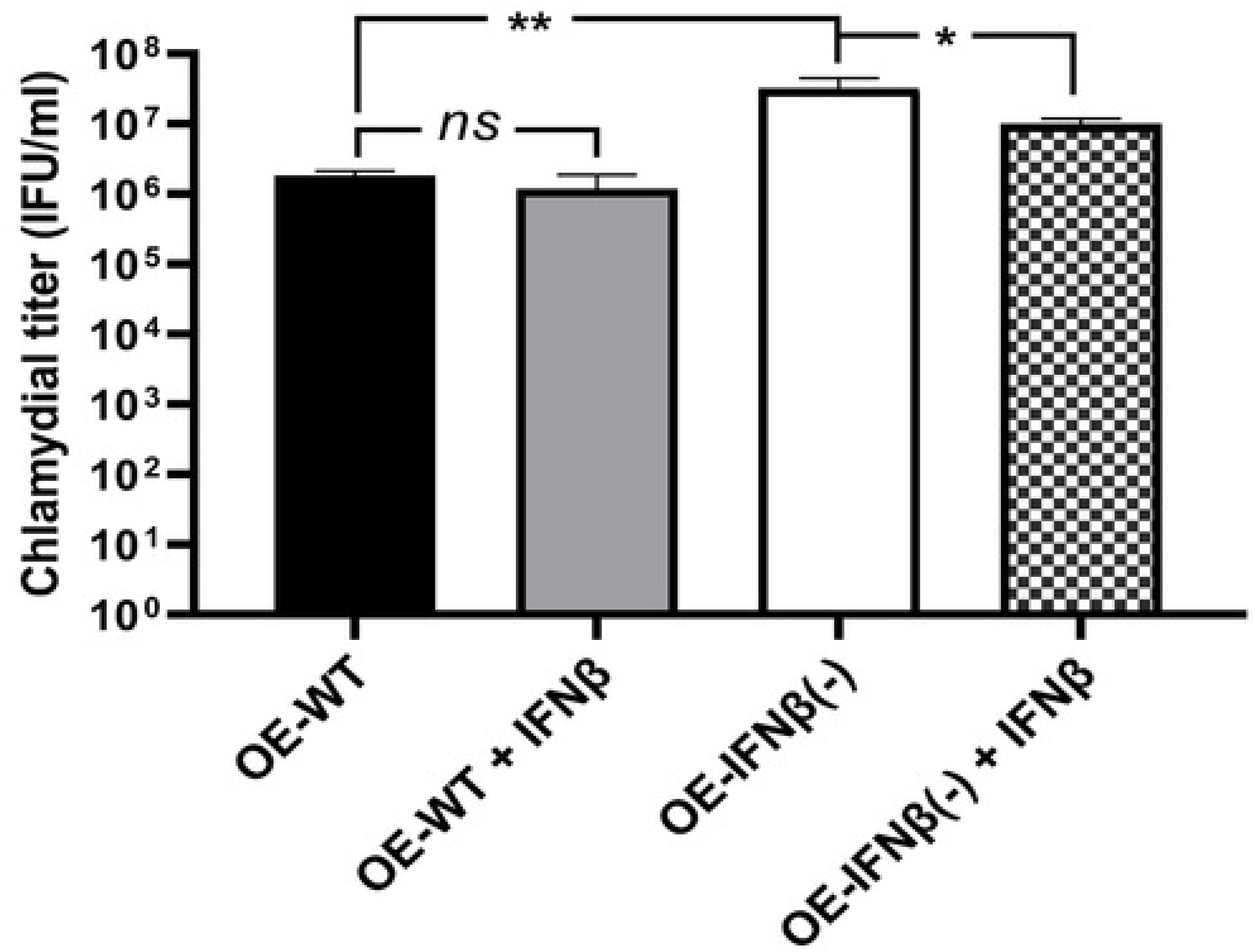
Impact of IFN-β deficiency on *C. muridarum* replication. OE-WT cells and OE-IFNβ (-) cells were infected with 5 IFU of *C. muridarum*/ cell in the presence or absence of 50U of recombinant IFN-β/ml. Lysates were collected, sonicated, and titrated on McCoy cell monolayers. The data presented are representative of three different experiments. *\*\*= p<0.01 *= p<0.05; ns= not statistically significant when compared as indicated*.

These findings demonstrate that epithelial-derived IFN-β directly restricts *Chlamydia* replication in OE cells and confirm that the enhanced bacterial burden observed in IFNβ–deficient cells is a direct consequence of IFN-β loss.

### IFN-β deficiency results in enhanced *Chlamydia muridarum* replication in vivo

To determine whether the epithelial-intrinsic effects of IFN-β observed in vitro translated to the *in vivo* setting, we next examined the course of genital tract infection in wild-type and IFNβ–deficient mice following intravaginal challenge with *C. muridarum*. Vaginal swabs were collected longitudinally, and chlamydial burden was quantified by enumerating recoverable IFU on McCoy cell monolayers.

IFNβ–deficient mice exhibited significantly increased *C. muridarum* shedding compared to wild-type mice over the course of infection (Fig. 8). Differences in bacterial burden were evident at multiple time points, indicating impaired control of infection in the absence of IFN-β. These findings mirror the enhanced intracellular replication observed in IFNβ–deficient OE cells and support a role for IFN-β in limiting *Chlamydia* replication within the genital tract.

**Fig 8.**
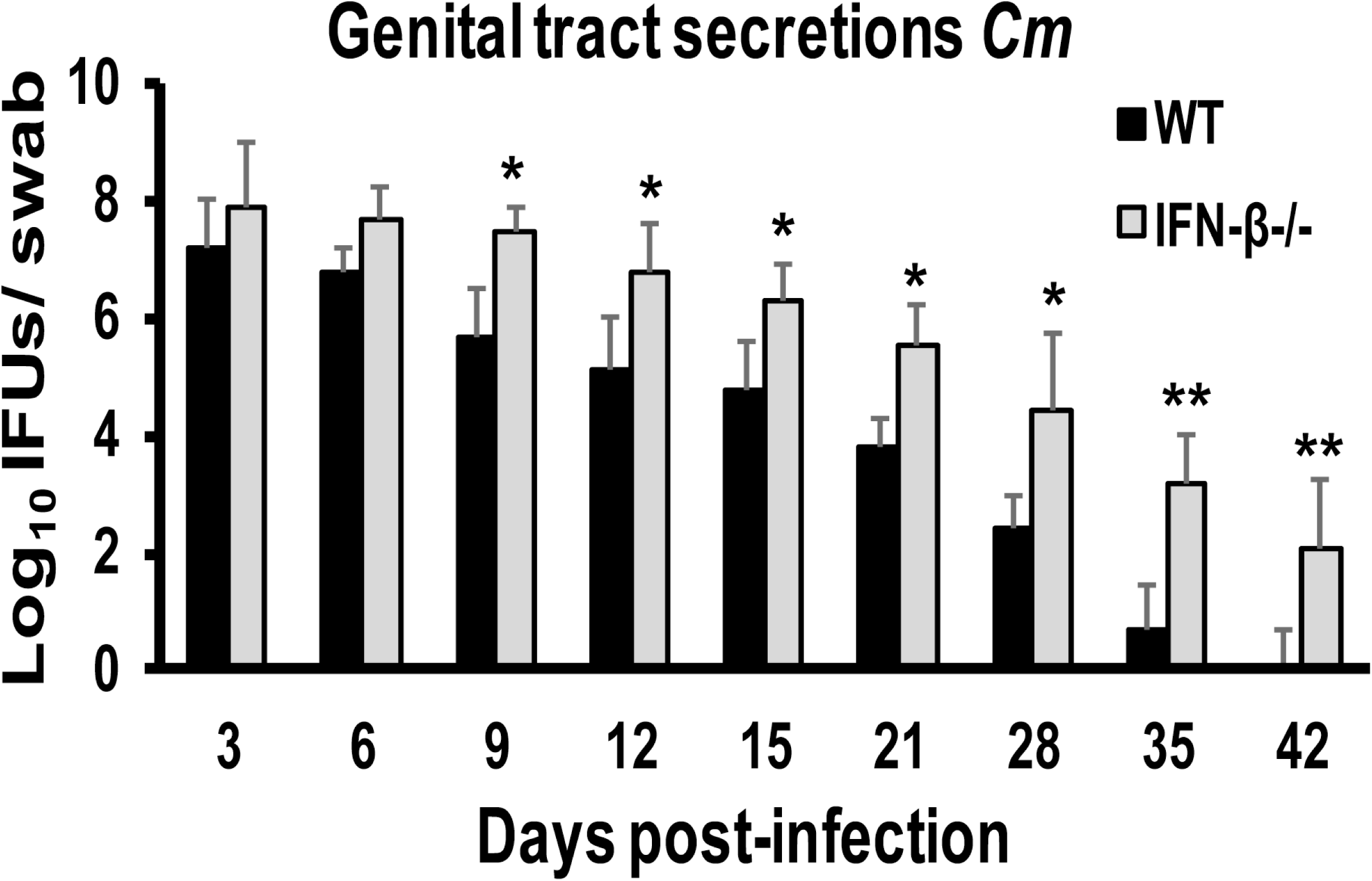
*C. muridarum* replication in the genital tracts of wild-type and IFNβ-deficient mice. Genital tract infections were performed in wild-type (WT) and IFNβ-deficient (IFN-β-/-) mice, and swabs were collected until day 42 as described in Materials and Methods. *C. muridarum* (Cm) titers were determined by infecting fresh McCoy cell monolayers. Combined data, n=18 mice for WT; n=17 mice for IFN-β^-/-^. IFU= inclusion forming units. *\*= p <0.05; **= p <0.005 when compared as indicated*.

Together, these data demonstrate that IFN-β contributes to host control of *Chlamydia* infection both at the epithelial cellular level and *in vivo*, and support a model in which IFN-β functions as a critical mediator of epithelial-driven host defense during genital tract infection.

## DISCUSSION

In this study, we demonstrate that interferon beta (IFN-β) plays a critical, non-redundant role in epithelial host defense during *Chlamydia muridarum* infection. Using well-characterized murine oviduct epithelial (OE) cell lines derived from primary oviduct epithelial cells and maintained under non-transformed conditions, we show that loss of IFN-β results in impaired induction of key inflammatory immune mediators, dysregulation of genes associated with immune regulation and fibrosis, altered chlamydial inclusion morphology, enhanced bacterial gene expression, and increased intracellular replication. Importantly, restoration of IFN-β signaling through exogenous supplementation rescues both epithelial immune responses and bacterial control, establishing a direct, cell-intrinsic role for IFN-β in limiting *Chlamydia* infection. These epithelial findings are supported by *in vivo* observations demonstrating enhanced bacterial burden in IFNβ-deficient mice.

The IFNβ-deficient OE cell lines used in this study are functionally equivalent during *C. muridarum* infection as other OE cells that were derived in our lab (3, 10). The OE cells used in this study were originally derived from epithelial cells isolated from the lumen of murine oviducts and have been extensively characterized as non-transformed epithelial models that retain innate immune responsiveness when cultured in keratinocyte growth factor (KGF)–supplemented media (10). As such, these cells provide a physiologically relevant system for interrogating epithelial-intrinsic immune mechanisms while avoiding confounding effects associated with oncogenic transformation or short-term primary cultures (22–24). The concordance between our *in vitro* findings and ongoing *in vivo* studies further supports the biological relevance of this epithelial model.

### IFN-β as a central epithelial regulator of host defense and bacterial development

Our data demonstrate that IFN-β contributes to epithelial host defense at multiple levels, including regulation of inflammatory mediator production, restriction of intracellular bacterial growth, and modulation of the chlamydial developmental cycle. The coordinated enhancement of chlamydial gene expression observed in IFNβ–deficient OE cells encompassed genes associated with bacterial viability (16S rRNA), metabolic activity (pyruvate kinase; PyK), structural components (outer membrane protein A; OmpA), and cell division (FtsK) (20, 25, 26). These genes were selected to span distinct stages of the chlamydial developmental cycle, thereby allowing assessment of whether IFN-β selectively impacts a single phase or broadly restricts bacterial development.

The observed upregulation of all four genes in the absence of IFN-β indicates that epithelial IFN-β does not merely delay clearance of *Chlamydia* but instead constrains multiple stages of intracellular growth and differentiation. This interpretation is further supported by the altered inclusion morphology observed in IFNβ-deficient OE cells and the increased recovery of infectious progeny. Together, these findings position IFN-β as a key epithelial-intrinsic regulator that limits *Chlamydia* replication by shaping the intracellular environment required for efficient bacterial development.

### Differential regulation of inflammatory mediators by IFN-β

Our analyses also revealed that IFN-β selectively regulates a subset of inflammatory immune mediators in OE cells during *Chlamydia* infection. The diminished expression of CCL5 (RANTES) and CXCL10 (IP-10) in IFNβ–deficient cells was anticipated, as both chemokines are well-established interferon-stimulated genes and have been repeatedly linked to IFN-β–dependent signaling pathways during intracellular infection (18). In contrast, the impaired induction of IL-6 and CXCL16 in the absence of IFN-β was unexpected, as these mediators are not classically regarded as IFN-β–dependent.

These findings suggest that IFN-β may influence epithelial inflammatory responses through both canonical interferon-stimulated gene pathways and less well-defined regulatory mechanisms. In particular, IFN-β may indirectly modulate IL-6 and CXCL16 expression by shaping the broader epithelial transcriptional landscape, influencing PRR crosstalk, or altering the balance of pro- and anti-inflammatory signaling during infection. The identification of IL-6 and CXCL16 as components of IFN-β–dependent epithelial responses expand the functional scope of IFN-β in the genital tract and highlights previously underappreciated links between interferon signaling and inflammatory mediator production during *Chlamydia* infection.

### Reconciling IFN-β–mediated protection with IFNAR-dependent pathology

A central implication of this study is the need to reconcile the protective role of IFN-β demonstrated here with prior reports indicating that global type I interferon signaling exacerbates pathology during genital tract *Chlamydia* infection. Studies employing interferon-α/β receptor–deficient (IFNAR^-/-^) mice have concluded that type I interferons promote inflammation and tissue damage, thereby reducing pathology in the absence of IFNAR signaling (11–13). However, IFNAR deficiency ablates signaling by all type I interferons, including multiple IFN-α subtypes and IFN-β, thereby precluding resolution of the individual contributions of these cytokines (27).

Our previous work provides a mechanistic framework for resolving this apparent paradox (28). In that study, we demonstrated that TLR3 deficiency during *Chlamydia* infection led to increased transcription of IFN-α2 and IFN-α4, whereas IFN-β expression exhibited distinct regulatory kinetics. Neutralization of IFNAR signaling effectively blocked late induction of IFN-α subtypes but had minimal impact on IFN-β transcription in OE cells, suggesting that IFN-β synthesis late during infection may be less dependent on canonical IFNAR-mediated autocrine signaling. These findings raised the possibility that IFN-β and IFN-α subtypes are differentially regulated and may exert divergent effects on host defense and pathology.

Building on this framework, the present study supports a model in which IFN-β functions as a protective, epithelial-intrinsic mediator that restricts *Chlamydia* growth, while excessive or dysregulated signaling by IFN-α subtypes (and potentially other type I interferons) may contribute to immunopathology. Under this model, the reduced pathology observed in IFNAR-deficient mice may reflect attenuation of pathogenic IFN-α-driven inflammatory pathways rather than loss of IFN-β–mediated host defense. Conversely, selective loss of IFN-β compromises epithelial control of bacterial replication, resulting in enhanced bacterial burden and downstream inflammatory consequences.

### Broader implications of type I interferon diversity in *Chlamydia* infection

While this study focuses on IFN-β and IFN-α as prototypical type I interferons, it is important to note that the type I interferon family encompasses multiple additional subtypes that may influence the outcome of *Chlamydia* infection (13, 29–31). Emerging evidence suggests that distinct type I interferons can exert non-overlapping, and in some cases opposing, effects depending on the tissue context, responding cell type, and timing of expression (32). Future studies examining the roles of individual IFN-α subtypes and other type I interferons will be necessary to fully define how this diverse cytokine family shapes genital tract immunity and pathology during *Chlamydia* infection.

### Integration with ongoing *in vivo* studies and future directions

The epithelial-intrinsic mechanisms defined here provide a strong foundation for understanding *Chlamydia* pathogenesis *in vivo*. Ongoing parallel studies examining genital tract infection in IFNβ–deficient mice demonstrate enhanced bacterial burden and immune dysregulation, confirming that the epithelial phenotypes observed *in vitro* translate to the intact host. These findings underscore the central role of epithelial IFN-β signaling in coordinating local immune responses that limit bacterial replication and tissue damage.

Based on the collective findings from this and prior studies, we propose that differential regulation of IFN-β and IFN-α signaling pathways represents a critical determinant of protective versus pathogenic immune responses during *Chlamydia* infection. Future investigations aimed at selectively dissecting IFNα-dependent pathways will be essential for clarifying their contributions to genital tract immunopathology and for identifying potential therapeutic targets that preserve host defense while minimizing tissue damage.

## Materials and Methods

### Ethics Statement

All animal experiments were conducted in accordance with protocols approved by the Indiana University Institutional Animal Care and Use Committee (IACUC) and were performed in compliance with NIH guidelines for the care and use of laboratory animals. All mice were monitored daily for lethargy, signs of vaginal bleeding, and/ or death. None of the mice exhibited morbidity during these studies. All mice were euthanized by either exposure to isoflurane or inhalation of carbon dioxide, followed by exsanguination.

### Mice

Wild-type C57BL/6 mice were purchased from The Jackson Laboratory (Bar Harbor, ME). and IFNβ–deficient mice generated on a C57BL/6 background (33) were graciously supplied by Michael S. Diamond (Washington University School of Medicine, Saint Louis, Missouri, USA) for use in the *in vivo* studies. Female mice aged 6-8 weeks were used for genital tract infection experiments. Animals were housed under specific pathogen-free conditions with *ad libitum* access to food and water and kept on a standard 12-hour light/dark cycle. All mice were given at least 1 week to acclimate to their new environment.

### Murine Oviduct Epithelial Cell Lines and Culture Conditions

Wild-type C57Bl/6 murine oviduct epithelial (OE) cell lines were derived from epithelial cells isolated from the lumen of mouse oviducts as previously described (10). Wild-type and TLR3-deficient OE cell lines were generated and characterized in prior studies (3). OE cell lines derived from IFNβ–deficient mice were generated for the current study using the same methodology.

All OE cell lines are non-transformed and retain innate immune responsiveness. Cells were maintained at 37°C in a humidified 5% CO₂ incubator in epithelial growth medium consisting of a 1:1 mixture of Dulbecco’s modified Eagle medium and Ham’s F12 medium (Sigma; St. Louis, MO) supplemented with 10% fetal bovine serum (FBS; HyClone), 2 mM L-alanyl-L-glutamine, bovine insulin (5 μg/ml), and recombinant human keratinocyte growth factor (KGF Sigma; 12.5 ng/ml) as previously described (3, 10).

### *Chlamydia muridarum* Propagation and Titration

*Mycoplasma*-free *Chlamydia muridarum* (strain Nigg) was propagated in McCoy cells as described previously (10, 34). Briefly, infected monolayers were harvested, and elementary bodies (EBs) were purified, resuspended in sucrose-phosphate-glutamate (SPG) buffer, and stored at −80°C until use. Infectious titers were determined by serial dilution and enumeration of inclusion-forming units (IFU) on McCoy cell monolayers. The total number of IFUs per ml was calculated by enumerating fluorescent inclusion bodies in infected McCoy cells, using an antibody specific for chlamydial LPS to detect them. Detection of the chlamydial LPS was performed using an Alexa Fluor 488-conjugated anti-mouse IgG secondary antibody (Invitrogen/Life Technologies; Carlsbad, CA), and immunostaining results were scanned and recorded using an EVOS imaging system (Thermo-Fisher, Pittsburgh, PA).

### *In vitro* Infection of Oviduct Epithelial Cells

OE cells were seeded into tissue culture plates and grown to confluence prior to infection in 6-, 12-, or 24-well plates. Cells were infected with *C. muridarum* at the indicated multiplicity of infection (MOI or IFU per cell) in culture medium as described previously (3, 14). Different MOIs were used depending on assay sensitivity, and mock-infected controls received an equivalent volume of SPG buffer. After centrifugation to synchronize infection, cultures were incubated at 37°C for the indicated times post-infection. Where indicated, recombinant murine IFN-β was added to cultures at a final concentration of 50 U/ml at the time of infection as described in (3, 15, 35). The media containing recombinant murine IFN-β remained on the cells throughout the infection. Untreated cells and SPG buffer-only treatment were used as uninfected and mock-infected controls, respectively. Supernatants and cell lysates were collected at specified time points for downstream analyses.

### Measurement of Cytokine and Chemokine Production

Culture supernatants were collected at the indicated time points post-infection and clarified by centrifugation. Levels of CCL5, CXCL10, IL-6, and CXCL16 were quantified using commercially available ELISA kits according to the manufacturer’s instructions (R&D Systems). Cytokine concentrations were calculated using standard curves generated with recombinant proteins.

### RNA Isolation and Quantitative Real-Time PCR

Total RNA was isolated from OE cells using the RNeasy kit plus (Qiagen, Valencia, CA). The DNA-free RNA was quantified using the NanoDrop spectrophotometer (Thermo Scientific), and cDNA was obtained with the Applied Biosystem’s high-capacity cDNA reverse transcription kit (Thermo Fisher).

Quantitative real-time PCR (qPCR) was performed using gene-specific primers for host immune mediators (including IL-10, MMP9, IL-4, TNF-α, CCL5, and IFN-γ) and chlamydial genes (16S rRNA, pyruvate kinase [PyK], outer membrane protein A [OmpA], and FtsK). Table 1 lists all target genes and primer sequences used for qPCR in this study. Relative gene expression was calculated using the ΔΔCt method with normalization to appropriate host or bacterial housekeeping genes. Relative expression levels were measured as a fold increase in mRNA expression versus mock controls and calculated using the formula 2^−ΔΔCt^ as described in (36).

**Table 1.**
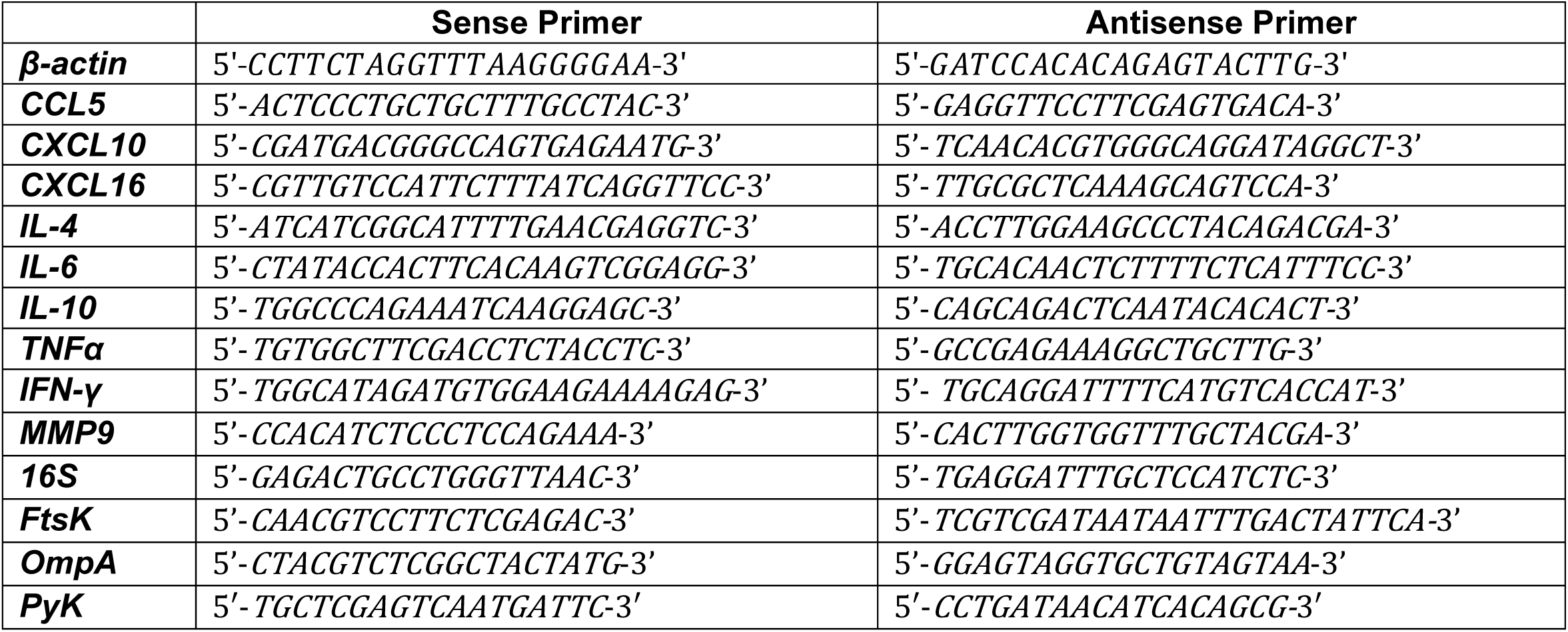
Primers for qPCR.

### Immunofluorescence Microscopy

OE cells were grown on glass coverslips and infected with *C. muridarum* as described above. At 24 hours post-infection, cells were fixed with paraformaldehyde, permeabilized, and stained with a *Chlamydia*-specific, fluorescein-conjugated monoclonal antibody (Fisher Scientific). Nuclei were counterstained with 4,6-diamidino-2-phenylindole (DAPI; Life Technologies) according to the manufacturer’s protocol and imaged at 40X under oil immersion using a Nikon Eclipse Ti microscope.

### *In vitro* Chlamydial Replication Assays

To quantify chlamydial replication, WT and IFNβ-deficient OE cells were plated in 24-well tissue culture plates and were either mock-infected, infected with 5 IFU/ml C. *muridarum*, or pre-treated with 50 U/ml recombinant IFN-β, then infected with 5 IFU/ml *C. muridarum* as described above. At 30 hours post-infection, cells were harvested by mechanical scraping with a pipette tip in 500μl of SPG buffer and frozen at −70°C until further processing. In order to study infectivity, the collected infected cell samples were vortexed, sonicated for 15 min in a water bath, and 50μl of the sample was passaged onto a fresh layer of McCoy cells for titration as described (10, 34) to determine recoverable IFU.

### *In vivo* Genital Tract Infection

Mouse infections were performed as described in (37), with minor modifications. Female mice were pretreated with 2.5 mg of Depo-Provera (medroxyprogesterone acetate; Pfizer, New York, NY) in 0.1 ml one week before vaginal infection with 10^5^ IFU *C. muridarum* (approximately 100 times the ID50) in 10µl of SPG (sucrose-phosphate-glutamic acid) buffer. Infection was monitored by swabbing the vaginal vault and cervix with a calcium alginate swab (Spectrum Medical Industries, Los Angeles, CA), and titers (IFU) of *C. muridarum* collected on the swabs were determined on McCoy cell monolayers as described (10, 34). Each mouse strain (wild-type and IFNβ-deficient) was infected in groups of five, and each experiment was repeated three times. To alleviate any possible distress, the mice were also briefly anesthetized with isoflurane prior to either intravaginal infection with *C. muridarum*, or the insertion of calcium alginate swabs. Swab samples were processed and titrated on McCoy cell monolayers as described above to determine bacterial burden over time.

### Statistical Analysis

Numerical data are presented as mean ± (SD). <u>All</u> experiments were repeated at least three times, and data were analyzed using appropriate statistical tests as indicated in the figure legends. Data were tested for normality prior to parametric analysis. Comparisons between groups were performed using Student’s *t* test or analysis of variance (ANOVA) in GraphPad Prism where appropriate. A *p*-value of <0.05 was considered statistically significant.

## Acknowledgements

This project was supported by National Institutes of Health Grants 1R01AI104944-01 and 1R21AI175847, and Marian University-Wood College of Osteopathic Medicine’s Office of Sponsored Programs and Research FRD grant funds. We graciously thank Michael S. Diamond at Washington University School of Medicine, Saint Louis, Missouri, USA for providing us with the use of the IFN-β KO mice used in these studies. We thank James Williams, Brahim Qadadri, and staff at the Indiana University infectious disease diagnostic laboratory for assistance with quantitative real-time PCR.

